# Unraveling eye movement-related eardrum oscillations (EMREOs): how saccade direction and tympanometric measurements relate to their amplitude and time course

**DOI:** 10.1101/2024.11.05.622042

**Authors:** Nancy Sotero Silva, Christoph Kayser, Felix Bröhl

## Abstract

Eye movement-related eardrum oscillations (EMREOs) reflect movements of the tympanic membrane that scale with the magnitude and direction of saccades. EMREOs have been consistently described in humans and non-human primates, yet many questions regarding this phenomenon remain unresolved. Based on bilateral in-ear recordings in human participants we here explore several properties of these EMREOs in order to improve our understanding of this signal’s origin and functional significance. Our data support that the EMREO time course is comparable between the left and right ears, and between paradigms guiding saccades by visual and auditory target stimuli. However, the precise amplitude time course differs significantly between ipsi-and contralateral saccades in addition to the previously known phase-inversion described for saccades in opposing directions. Finally, our data suggest that the EMREO amplitude is negatively related to the compliance of the tympanic membrane as established by tympanometry. Collectively, these results support the notion that EMREOs reflect motor-related top-down signals relayed to the ear from yet-to-be-resolved sources, and fuel the speculation that EMREOs may be generated by the middle ear muscles in a differential operation similar to the execution of ipsi-and contralateral saccades.

## 1. Introduction

In-ear microphone recordings reveal rhythmic signals that are systematically aligned to saccadic eye movements and which last for up to 100 ms (Gruters et al., 2018; Bröhl and Kayser, 2023; Lovich et al., 2023a). These so-called eye movement-related eardrum oscillations (EMREOs) have been described in humans and non-human primates, and reflect movements of the tympanic membrane that scale with the magnitude and direction of saccades. For instance, saccades towards locations ipsi-and contralateral to the ear in focus induce signals of opposing phase, which has fueled speculations of whether EMREOs may reflect the spatially-specific alignment of ear-and eye-centered signals (Cho, Ravicz and Puria, 2023; Lovich et al., 2023a; Tasko et al., 2022; Caruso et al., 2021; Razavi, O’Neill and Page, 2007). Yet, many questions regarding the phenomenology, origin, and functional role of EMREOs remain unresolved (King et al, 2023; Lovich et al., 2023b). We here focus on four specific questions described in the following.

First, current evidence suggests that EMREOs are generated by active processes in the cochlea or middle ear. One possibility is that the two middle ear muscles (MEMs), i.e. the stapedius and tensor tympani muscles, induce the EMREOs (Gallagher, Diop and Olson, 2021). These muscles connect to the tympanic-ossicular system and adjust sound transmission in loud environments, such as during the acoustic reflex (Cho, Ravicz and Puria, 2023; Lovich et al., 2023b; Edmonson et al., 2022; Tasko et al., 2022). The stapedius and tensor tympani muscles are innervated by different cranial nerves (the seventh and the fifth, respectively) and the respective acoustic reflexes involve partly distinct circuits. The pathways activating the respective motor neurons involve structures that receive multisensory and motor-related signals, whose origin remains in part unknown (Mukerji, Windsor and Lee, 2010; Borg, Counter and Rösler, 1984; Møller, 1984). Intriguingly, movements of the eyes towards ipsi-and contralateral directions involve opposing muscles, which are innervated by distinct cranial nerves - the lateral rectus muscle being controlled by the sixth and the medial rectus muscle by the third cranial nerve (Pierrot-Deseilligny, 2011; Machado, 2004; Walker, Hall and Hurst, 1990). It is therefore conceivable that the middle ear muscles could be governed by a similar principle, in that information about opposing saccade directions enters the pathways towards the stapedius and tensor tympani muscles in a differential manner. Indeed, previous studies have shown that the sign of the EMREO deflections differ between saccades in opposing direction, but also noted that the precise amplitude time course may possibly differ between ipsi-and contralateral saccades beyond the opposing sign of the respective signals (Gruters et al., 2018; Murphy et al., 2020; Lovich et al., 2023a). To substantiate this observation, we aimed for a systematic and statistical comparison of potential differences in the EMREO amplitude time course between saccade directions. To this end we performed bilateral in-ear recordings in human volunteers and compared the EMREO amplitude time course between horizontal saccades to the left and right.

Second, previous studies on otoacoustic emissions have noted potential differences between such emissions recorded from left and right ears (Keefe et al., 2008; Sininger and Cone-Wesson, 2004). While EMREOs and otoacoustic emissions are likely generated by different mechanisms, it remains possible that the precise EMREO time course differs systematically between ears, potentially reflecting other patterns of hemispheric specialization in the auditory system (Zatorre, 2022; Jamison et al., 2006; Schönwiesner, Rübsamen and von Cramon, 2005). Hence, we systematically compared the EMREO time course between left and right ears.

Third, the pathways towards the middle ear motor neurons may possibly involve both descending sensory information and descending motor-related signals. Therefore, it remains unclear whether the EMREO time course is shaped solely by the motor-related properties of the eye movement or also by the nature of the external sensory signal guiding a saccade. Previous work has shown that EMREOs emerge in relation to visually-guided saccades, during free-viewing of images, and in darkness (Gruters et al., 2018; King et al., 2023; Lovich et al., 2023a, 2023b). If EMREOs are directly and only driven by a motor-related signal, one may not expect differences between EMREOs induced by saccades made in response to target stimuli presented in different sensory modalities, such as visual and auditory saccade targets. However, task-guided and spontaneous saccades may also differ by the contributing cerebral pathways (Bon and Lucchetti, 1988; Lucchetti and Bon, 1997; Leigh and Kennard, 2004). We here address this question by contrasting EMREOs induced by visually and auditory guided saccades.

Lastly, we studied whether individual variations in the EMREO covary with hearing status and middle ear function. Assuming that EMREOs are generated by the MEMs, it is possible that the amplitude of the EMREO varies across individuals with the mobility or functioning of the tympanic membrane. One possibility is that a reduced compliance of the tympanic membrane to changes in air pressure results in a reduced EMREO amplitude. Indeed, previous work already speculated that the resonance properties of the middle ear structures might be linked to the variability in the EMREO signal (Lovich et al. 2023a). We here directly tested for a relation between the EMREO amplitude and middle ear properties by systematically characterizing participants’ middle ear function and hearing status using tympanometry, measurement of acoustic reflexes and audiometric testing.

Our results show that EMREOs are comparable between the left and right ears, are comparable when guided by visual or auditory targets, but differ in amplitude time course between saccades towards targets ipsi-and contralateral to the respective ear. Furthermore, the amplitude of the early phase of an individual EMREO is negatively related to the compliance of the tympanic membrane as obtained by tympanometry.

## 2. Materials and Methods

### 2.1. Participants

Overall, we collected data from 34 participants (24 females, 10 males, mean ± SD age = 27,3 ± 5,3 years) recruited among university students and community members. They participated following written informed consent and received 15 euros per hour as compensation for their time. The study was approved by the ethics committee of Bielefeld University (#2021-218). Participants were screened and selected as follows: they should not report known hearing or vision impairments (vision corrected to normal with glasses or lenses < ± 2 dpt was accepted) and no known neurological or psychiatric disorders. We visually inspected their external ear canals and performed pure tone audiometry to establish air conduction hearing thresholds (MADSEN Xeta, GN Otometrics at 250, 500, 1000, 2000 and 4000 Hz). We also performed tympanometry (R26M Middle Ear Analyzer, Resonance Audiology, 226 Hz probe, sweeps from 200 to-300 daPa) and tested for acoustic reflexes at 500, 1000, 2000 and 4000 Hz. Ipsi-and contralateral reflexes were obtained based on the equipment’s “Automatic” setting: stimuli were presented for 1 s in 5 dB steps, starting at 80 dB until the intensity that provoked a 0.05 mmho variance (indicating the trigger of a reflex), or until a fixed threshold was reached; these were 100 dB (500 and 4000 Hz) and 110 dB (1000 and 2000 Hz) for ipsilateral stimulation, and 120 dB for contralateral stimulation. Individuals with a pure tone audiometry (PTA) threshold average below 20 dB were considered to have normal hearing (World Health Organization, 2020). Six participants did not complete the full experiment due to either visible ear obstructions (one participant), low performance in the task-related training blocks (two participants, see 2.3.), PTA thresholds above 20 dB (one participant) or voluntary withdrawal (two participants). For each participant, data was obtained in two sessions taking place on two different days. Based on the hearing screening, tympanometry results (i.e. tympanometric curve classified accordingly to the compliance peak values) and the equivalent ear canal volume, we ensured that the functioning of the ear was comparable between sessions. In each session we measured EMREOs in response to visually guided and auditory guided saccades, as explained below. As part of the experiments we also obtained data for protocols involving additional tasks, which are not reported here.

### 2.2. Experimental setup and data acquisition

The experiments were carried out in a soundproofed and electrically shielded booth with low ambient light levels. Participants were seated 100 cm away from a sound-permeable screen (Screen International Modigliani, 2 × 1 m) with their head on a chinrest aligned to the center of the screen. Visual stimuli were projected (Acer Predator Z650, Acer; 60 Hz refresh rate) onto this screen. Acoustic stimuli were presented from an array of 5 free-field speakers located behind the screen (Monacor MKS-26/SW, MONACOR) positioned evenly spaced between-9 and 9 degrees azimuth. Sounds were generated from a Sound BlasterZ Soundcard (sampling rate 44100 Hz) and amplified with a HeadAMP 4 (ARTcessories) amplifier. Stimuli were presented using the Psychophysics toolbox (Brainard, 1997) for MATLAB (The MathWorks, 2022), which was synchronized to the recording system using TTL pulses.

In-ear microphone recordings were performed using two Etymotic ER-10C (Etymotic Research) systems. Microphone signals were pre-amplified using the ER-10C DPOAE amplifiers (Etymotic Research), with gain set to +40 dB, and digitized through an ActiveTwo AD-box (BioSemi) at a sampling rate of 2048 Hz.

Eye movements were recorded from the left eye using the EyeLink 1000 plus eye-tracking system (SR Research) with a sampling rate of 1000 Hz. Eye-tracking calibration was performed at the beginning of each block using a 5-point grid. The parameters for saccade detection in the EyeLink system (“cognitive” setting) were a velocity threshold of 30°/s and an acceleration threshold of 8000°/s.

To monitor the quality of the microphone functioning and of the in-ear recordings, control measurements were done before the beginning of each session. These comprised an empty-room measurement consisting of 15 seconds of silence, then the presentation of 20 tone stimuli presented from the speakers (10 stimuli at 100 Hz, 10 stimuli at 200 Hz, duration 400 ms). The same measurement was repeated with the ear-probes inserted into the participants’ ear canals. The baseline noise spectrum and the signal amplitudes in response to these tones served as a parameter for adjustments to the microphone insertion. Furthermore, based on the empty-room recordings we observed a consistent difference in output levels of the left and right microphones for the probe tones of a factor of about 1.21. We used this factor to correct for the expected EMREO levels between ears in the subsequent analyses, thereby calibrating the data analysis for slight differences in the effective gain of the recording equipment.

### 2.3. Experimental design

We obtained data for three paradigms presented in different blocks in each session. First, we collected EMREOs in response to visually-guided saccades, similar to our previous work (Bröhl and Kayser, 2023). As illustrated in Figure 1, a white dot (0.2 degree radius, on the gray background of the screen) was presented at a position of 0 degrees azimuth for a fixation period of between 900 and 1400 ms and then jumped to a random position to the left or to the right. For the first session, target positions were between-9 and-3 degrees azimuth (left, in 1 degree steps) and 3 and 9 degrees (right, in 1 degree steps). For the second session we extended the spatial range of saccades to between-18 and - 6 degrees (2 degrees steps), 6 and 18 degrees (2 degrees steps). Thereby we ensured that the collective data samples a large range of saccade parameters. Participants were instructed to fixate their gaze on the dot and to follow its movement to the target position. Inter-trial intervals lasted between 1900 and 2400 ms. In each session participants performed two blocks of 140 trials each, with the directions and amplitudes of target jumps pseudo-randomized across trials.

**Figure 1.**
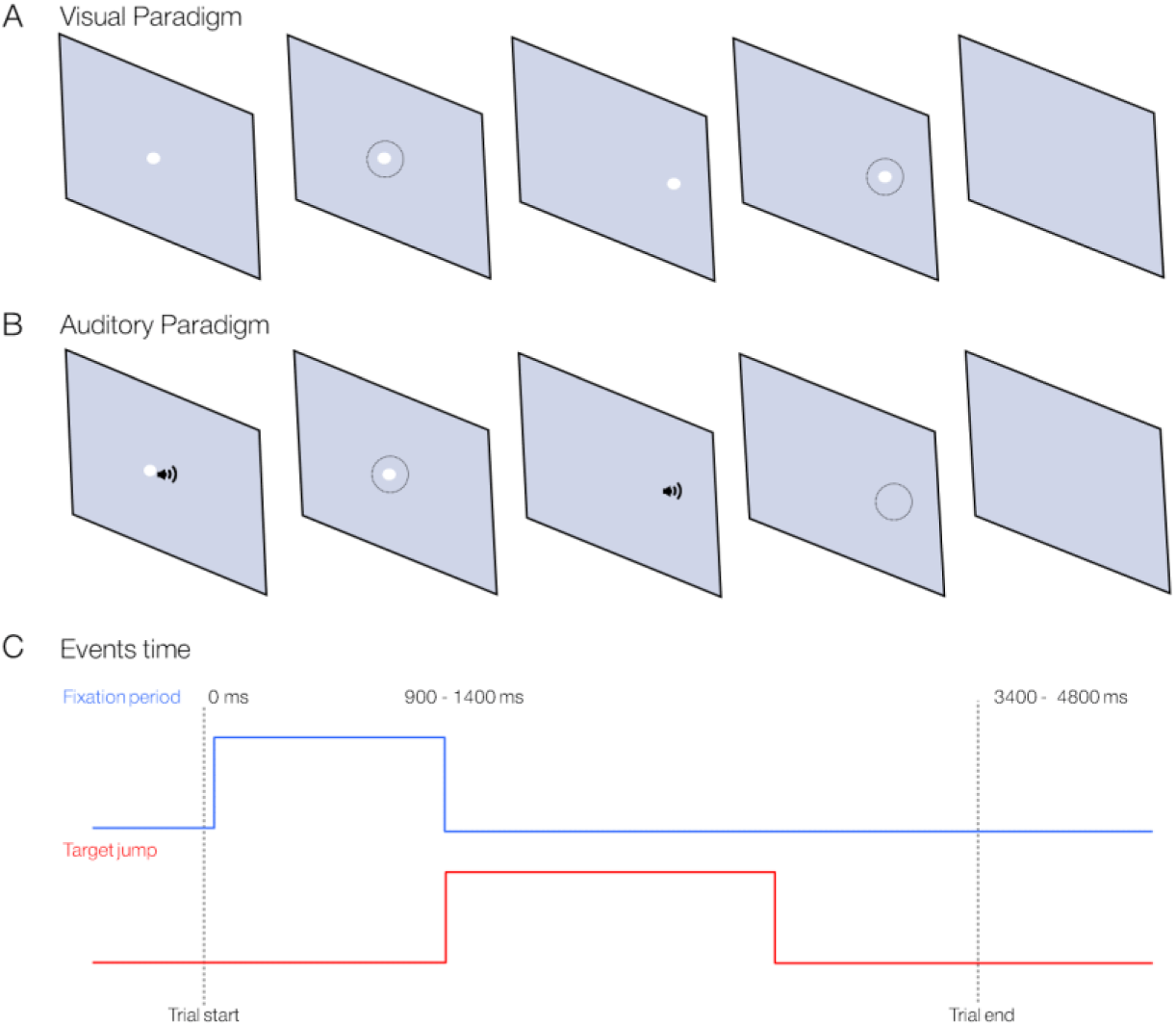
Schematic representation of the experiment setup. The figure shows the target sequence for the visual paradigm (**A**) and the auditory paradigm (**B**). The dots represent the central fixation position and the saccade targets in the visual paradigm, the loudspeaker symbols represent the horizontal positions of the tones that served as saccade targets in the auditory paradigm, and the dotted circle indicates the gaze position. Participants were instructed to fixate their gaze on the central position as long as the dot remained on screen. Subsequently, the central dot would disappear and either jump to one of the target positions in the visual-guided paradigm, or sounds were presented from the respective target location in the auditory-guided paradigm. Participants then performed a saccade towards the target and had to keep the gaze fixed until the end of the trial. Panel (**C**) represents the time distribution within one trial (the gray vertical line represents the trial limits). Fixation period would last for a randomly generated interval between 900 and 1400 ms, represented by the blue line. The saccade would be detected in the 1000 ms interval after the start of the target jump, represented by the red line.

Second, we collected EMREOs in response to auditory-guided saccades. To this end, two auditory target sounds were presented in each trial (clicks of 15 ms duration, smoothed by a 12.5 ms cosine ramp; around 80 dB SPL) from the speakers behind the screen. The first sound was presented at 0 degrees azimuth, while the second was presented 650 ms later. For the first session saccade targets jumped either to the left (at-9,-6 or-3 degrees azimuth) or to the right (at 3, 6 or 9 degrees), in the second session to-18°,-12° and 6° to the left or 6°, 12° and 18° to the right. Sound locations in between two speaker positions were created by presenting sounds from both speakers with linearly interpolated amplitudes. A central visual fixation was presented for 150 ms with the onset of the first tone to ensure central initial fixation, but disappeared before presentation of the second tone presenting the saccade target. Participants were told that the sound would only move horizontally and were instructed to make the saccade to the point on the screen matching the location of the perceived second sound. Participants performed two blocks of 90 trials each, with the directions and amplitudes of target jumps pseudo-randomized. Lastly, participants performed four blocks comprising visually-guided saccades in conjunction with an additional sensory task, the data of which are not reported here.

### 2.4. Data analysis

#### 2.4.1. Preprocessing

To extract EMREOs, the microphone recordings were epoched and aligned to the onset of the task-guided saccades as detected by the EyeLink system. For each paradigm and trial, we extracted the visually (or acoustically) task-guided saccade as the largest saccade reported by the eye tracking system in the 2.5 s period following the jump of the saccade target. We imposed the following requirements for saccades to be included in the analysis: (1) that the saccade amplitude was within the range 2 to 16 degrees - this was implemented mostly to avoid small saccades; (2) that the initial fixation prior to the saccade was stable - for this we required that the horizontal eye position was within a window of ± 4 SDs of the average fixation position across all epochs for a given participant and paradigm; (3) that the entire horizontal and vertical traces of the eye position were contained in an 18 x 18 degree window around the central fixation point (as outside this range eye tracking was unstable); (4) that the entire saccade was contained in the 2.5 s time window. The existence of a saccade with the required amplitude was the criterion that led to the exclusion of most trials, while few trials were rejected based on criteria 2 - 4. Although the saccade targets were set to visual angles of up to 18° in the second session, we noticed during the analysis stage that eye movements to these positions were sometimes not well tracked by the eye tracker. Hence, we excluded eye positions at these large eccentricities from further analysis. For the visual paradigm these criteria collectively led to the exclusion of around 15% of trials, while for the auditory paradigm these criteria led to the exclusion of around 68% of trials. Notably, for the auditory paradigm many trials did not contain a saccade of the required amplitude within the required time window, suggesting that participants had difficulties following the auditory targets within the given time frame of each trial.

The EMREO epochs for the included saccades were screened for extreme values, and epochs for which the maximal Z-score of the EMREO signal at any point in time exceeded 4.5 SDs of the overall signal were removed. This was the case for around 16% of epochs in the visual paradigm and 15% in the auditory paradigm. Given that the EMREO is best visible in the average across many repetitions of the same condition, we sought to maximize the amount of useful data for these analyses and collapsed the data from the same paradigm obtained in each session.

Based on the above criteria the data of another 5 participants had to be excluded, mostly due to excessively noisy in ear recordings or low quality of the eye tracking data. Overall, we report data from 23 participants. For these we retained 410 ± 10 (mean ± SEM) epochs per participant in the visual paradigm, corresponding to 73 ± 2% of the actually recorded trials; for the auditory paradigm we retained 98 ± 8 epochs, corresponding to 27 ± 2% of the recorded trials.

#### 2.4.2. EMREO Analysis

The main focus was to compare the deflections of the tympanic membrane i.e., the EMREO amplitude time courses between ears, between saccades to ipsi-and contralateral targets, and between visually and auditory-guided saccades. We implemented this comparison for each participant based on the epoch-averaged EMREO signals at individual time points around saccade onset. We opted for such a direct comparison of the amplitude time course of two signals, rather than e.g. a two-step approach based on regression models (Gruters et al., 2018; King et al., 2023; Lovich et al., 2023a) for the following reason. By comparing amplitude time courses, the statistical test directly focuses on the actual data of interest, similar e.g. as when comparing evoked responses in the EEG or local field potentials in in vivo data. An alternative approach, such as using regression models to describe the dependency of the EMREO time course on saccade properties (Murphy et al., 2020; King et al., 2023; Lovich et al., 2023b) and then testing for differences in the slope of specific regression predictors between conditions, is less direct and may add additional uncertainties. In fact, such an approach would be associated with more statistical uncertainty, as the regression model only provides an approximate descriptor of the EMREO signal.

To compare EMREOs between ears, or between ipsi-and contralateral saccades, we focused only on the saccades obtained in the visual paradigm. To compare visual and auditory guided EMREOs, we contrasted EMREOs between paradigms. In this analysis we also collapsed epochs across ears for each saccade direction, as the comparison of ears had revealed no significant differences (see session 3). To relate the EMREOs to the results from the hearing screening and tympanometry, we calculated for each participant and ear the EMREO amplitude as the sum of the total EMREO deflection (i.e. the absolute values) in a time window from 0 ms to 30 ms based on the data in the visually guided saccade paradigm. For comparison, we repeated this analysis also based on the EMREO amplitude in a later time window (30 ms to 80 ms). These windows were established based on the participant, ear and condition averaged EMREO time course, which featured prominent zero crossings near 30 and 80 ms (c.f. Fig. 4). We refer to these two time windows as the “early” and “later” EMREO phases. Changing these windows slightly did not lead to different overall conclusions.

#### 2.4.3. Analysis of eye tracking data

Although for a given trial we instructed participants to perform a horizontal saccade in a specific direction at a specific amplitude, the actual saccades may not perfectly follow this instruction, as they may also move the eyes along the vertical dimension, and could in theory even be executed in the opposite direction. We therefore verified that most saccades were performed in the appropriate direction. For this we computed the percentage of trials in which the horizontal saccade component was directed in the same direction as the target jump, and we also calculated the angular deviation from a straight horizontal line for each saccade.

From the eye tracking data, we extracted the horizontal and vertical eye traces, the time course of the overall velocity and the amplitudes, durations and peak velocities of each individual saccade. These saccade characteristics are shown in Figure 2 and illustrate that the saccades were comparable for eye movements to the left and right and between visual and auditory paradigms. To compare saccade properties between two conditions, we relied on non-parametric two-sample rank-sum tests after collapsing amplitudes (durations) across participants.

**Figure 2.**
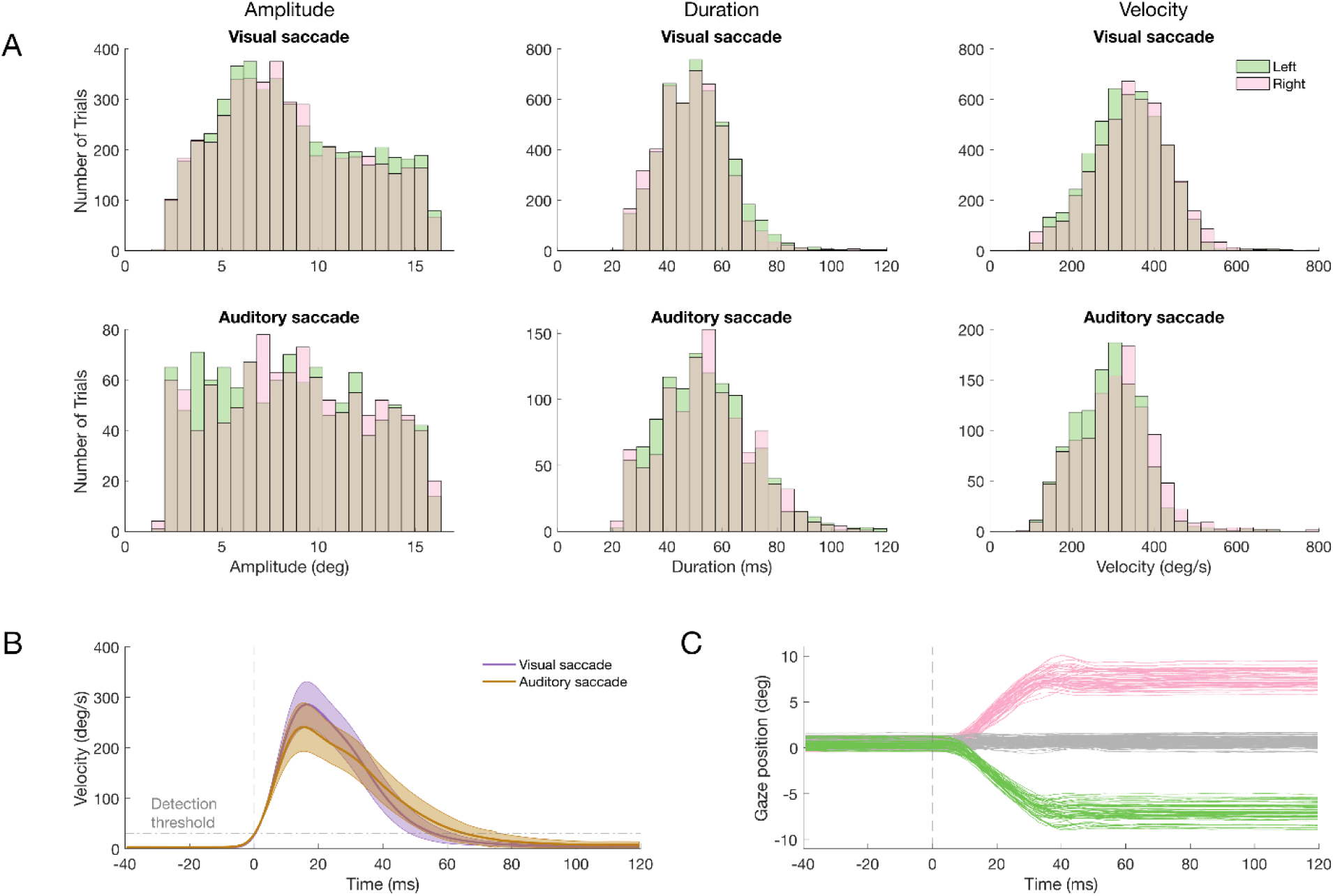
Properties of saccadic eye movements. (**A**) Distribution of amplitude, duration and peak velocity for saccades to the left and right directions in the visual (top row) and auditory (bottom row) paradigms. (**B**) Group average time courses for saccade velocities in the visual and auditory paradigms. The horizontal dashed line indicates the velocity detection threshold of the eye tracking system and the vertical dashed line indicates the respective saccade onset. Thick lines indicate the mean and shaded areas the standard error of mean across participants. (**C**) Saccade eye traces for one example participant in a one block, displaying momentary horizontal gaze positions for the saccades to left (green) and right (pink) targets, and the vertical gaze position (grey). Each line represents an individual trial.

#### 2.4.4. Statistical analysis

Group-level inference on the difference in the EMREO amplitude time course between conditions (ears, saccade directions or paradigms) was implemented using a cluster-based permutation test controlling for multiple comparisons along time (Nichols and Holmes, 2002; Maris and Oostenveld, 2007). The first level statistics was based on paired t-tests at each time point comparing the two conditions of interest within each participant. Time points with t-values reaching a first-level threshold of p < 0.01 were included in a clustering procedure, which were defined to require a minimal cluster-size corresponding to 7 ms. The maximum sum of each potential cluster was compared against a surrogate distribution, which was obtained by randomly flipping the sign of the condition-difference across participants for 10,000 times and recalculating the test statistics. We report clusters at a second-level significance of p < 0.05. For each cluster we report Cohen’s D as a measure of effect size at the time point of maximal difference. Correlations between hearing screening and EMREO amplitude were computed across ears (hence a sample size of 23 * 2) based on the Spearman’s rank correlation coefficient.

## 3. Results

### 3.1. Eye movements

We analyzed bilateral EMREO recordings obtained from 23 volunteers performing horizontal saccades in two paradigms, guiding these either by visual or auditory saccade targets (see Fig. 1). In the following we first report the properties and summary statistics of the respective eye movements before focusing on the EMREOs.

The experimental paradigms required participants to perform horizontal saccades in a given direction. We verified that for most trials the executed saccades were indeed in the correct (and not the opposite) horizontal direction. For the visual paradigm this was the case in 96.6 ± 2.7% (mean ± SEM across participants) of trials, for the auditory paradigm this was the case for 83.4 ± 2.3% of trials. We also verified that most saccades were made along the horizontal direction and did not contain major gaze changes along the vertical direction: the average absolute amplitude of the vertical gaze change was small (0.38 ± 0.02 degree for visual saccades, 0.82 ± 0.1 degree for auditory saccades) and much smaller than the minimal amplitude of the horizontal component (required to be at least 2 degrees). We also calculated the angular deviation from a purely horizontal saccade (regardless of the direction of deviation from the horizontal line), which was also comparably small and amounted to 2.7 ± 0.16 degrees angle for visual saccades and 6.8 ± 1.0 degrees angle for auditory saccades.

Figure 2A shows the main saccade characteristics, including amplitude, duration and peak velocity for the main comparisons of interest: saccades to the left and right in the visual paradigm and between visual and auditory paradigms. Statistical testing showed that for the visual paradigm saccade amplitudes did not differ significantly between movements to the left and right (rank-sum tests, p = 0.33, Z = 0.9), while saccades to the left were slightly longer compared to those to the right (median values 47 vs. 49 ms; p < 10-5, Z = 4.9) and featured higher peak velocities (334 vs 328 deg/s; p < 10-5, Z = 5.6). Comparing the visual and auditory paradigms we found no significant difference in saccade amplitudes (p = 0.76 Z = 0.3), while auditory guided saccades were longer than visual guided saccades (median values 53 ms vs 49 ms; p < 10-, Z =-11.5) and had lower peak velocities (300 deg/s vs. 341 deg/s, p < 10-5, Z =-19.2). Overall this suggests that the saccade amplitudes were comparable between the relevant conditions compared below. This is important, given that the saccade amplitude is predictive of the EMREO amplitude and differences in saccade amplitudes may otherwise confound the following comparison of EMREOs. While saccade durations differed between the conditions of interest, we note that these differences are numerically small and much shorter than the temporal structure of the EMREOs reported below.

Figure 2B illustrates the time course of saccade velocity relative to the detected saccade onset, which served as the alignment point of EMREO epochs. While the average velocity starts to increase prior to time point 0 s, the example eye traces in Figure 2C show that visible deflections of the direction of gaze emerge only a few milliseconds after 0, and hence sizable movements of the eye occur only subsequent to the alignment point of EMREOs.

### 3.2. The EMREO time course differs between ipsi-and contralateral saccades

Previous studies have shown that the sign of the main deflections in the EMREO differs between saccades towards opposing directions (Gruters et al., 2018; Lovich et al., 2023a; King et al., 2023; Bröhl and Kayser, 2023). Our data confirm this difference in EMREO sign. Figure 3A shows the group-level EMREOs for each ear and direction in the visual paradigm, revealing initial negative (positive) deflections for ipsi-(contra) lateral saccades around 10 to 15 ms, and an opposite pattern for the deflection around 25 ms. As the figure shows, even when adjusting for this difference in sign the precise amplitude time courses for ipsi-and contralateral saccades differ: in Fig. 3A the solid blue line shows the actual ipsilateral EMREO, while the dashed orange line shows the sing-inverted contralateral EMREO. This difference in time courses is shown directly in the panels below (Figure 3A, black lines).

**Figure 3.**
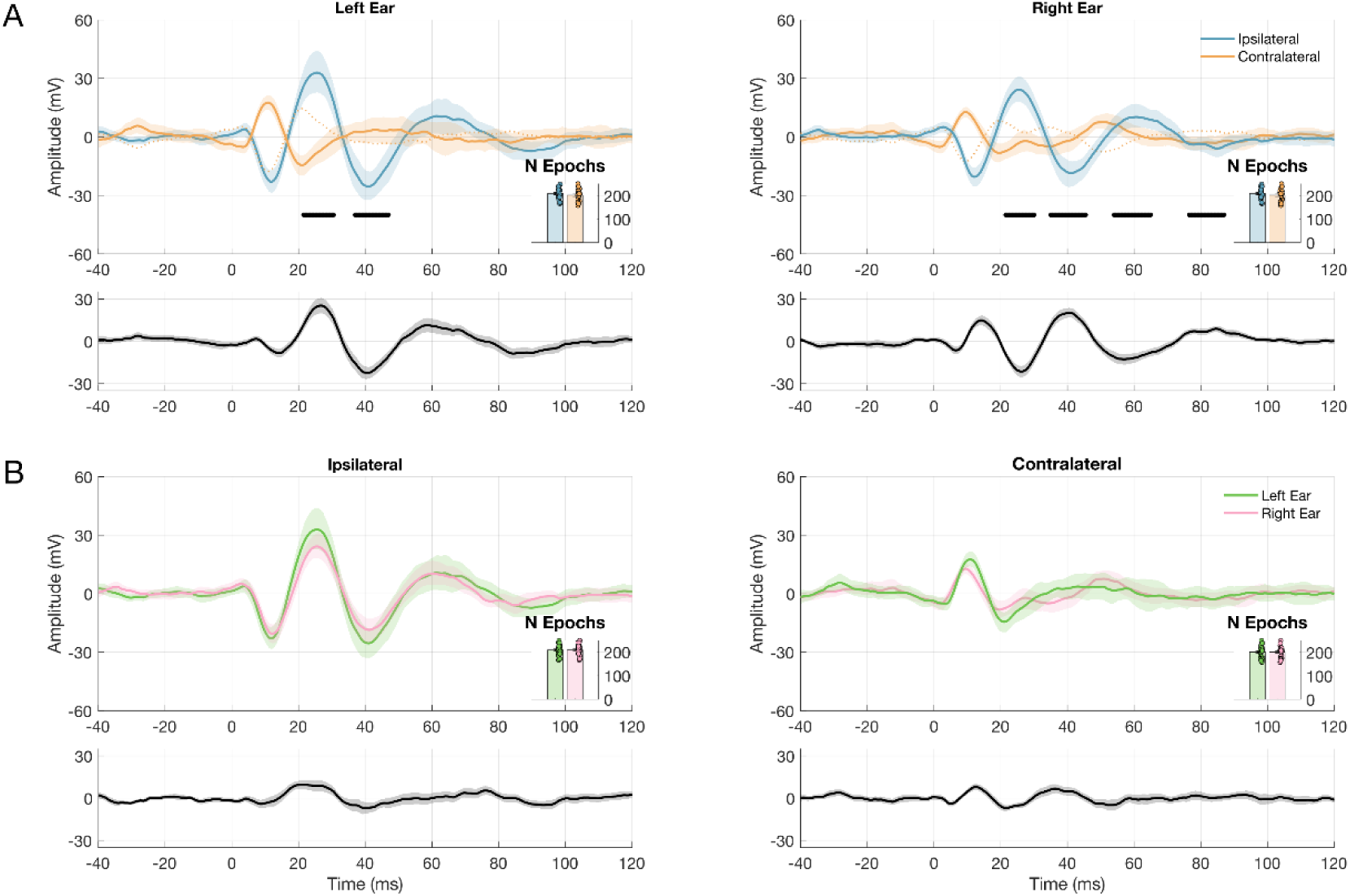
Group-level EMREOs for ipsi-and contralateral saccades in the visual paradigm. (**A**) The upper panels show the EMREO time courses for each ear and saccade direction (solid lines); the dashed lines show the ipsilateral data after inverting the sign. The lower panels show the difference between ipsi-and contralateral directions, after sign-inverting the ipsilateral data. (**B**) Comparison of EMREOs between ears for each saccade direction. Lines indicate the group mean (n = 23), shaded areas the 95th percentile bootstrap confidence interval. Black bars indicate the clusters reaching significance (at p <0.05). Insets indicate the number of data epochs per each participant and group.

We systematically tested for a difference in time courses of ipsi-and contralateral EMREOs after correcting for this difference in sign, separately for each ear using cluster-based permutation tests. These revealed significant differences for both ears (Left: positive cluster from 21.5 - 30.5 ms, Cohen’s D = 0.97, p = 0.001; negative cluster from 37 - 47 ms, Cohen’s D = 0.59, p = 0.0001; Right: positive cluster from 35 - 45.5 ms, Cohen’s D = 1.23, p = 0.0001; second positive cluster from 76.5 - 87 ms, Cohen’s D = 0.85, p = 0.002; negative cluster from 21.5 - 30 ms, Cohen’s D = 0.60, p = 0.001; second negative cluster from 54 - 65ms, Cohen’s D = 0.60, p = 0.002). Although the timing of significant differences does not allow inference about the precise time of an effect, these multiple and broad significant clusters corroborate that ipsi-and contralateral EMREOs differ not only in phase but also in the specific amplitude time course.

### 3.2. The EMREO time course does not differ between ears

Given that we systematically recorded EMREOs in both ears of each participant, we could directly compare the time courses between these. Figure 3B shows the direct comparison between ears for each saccade direction separately. A statistical comparison revealed no significant differences between ears (at p < 0.05; cluster-based permutation statistics) confirming that there is no evidence for a consistent difference in EMREO time courses between ears. Figure 4 shows the individual participant-wise EMREO traces for each ear, and illustrates the largely consistent timing of EMREO peaks during the first about 30 ms of the epoch.

**Figure 4.**
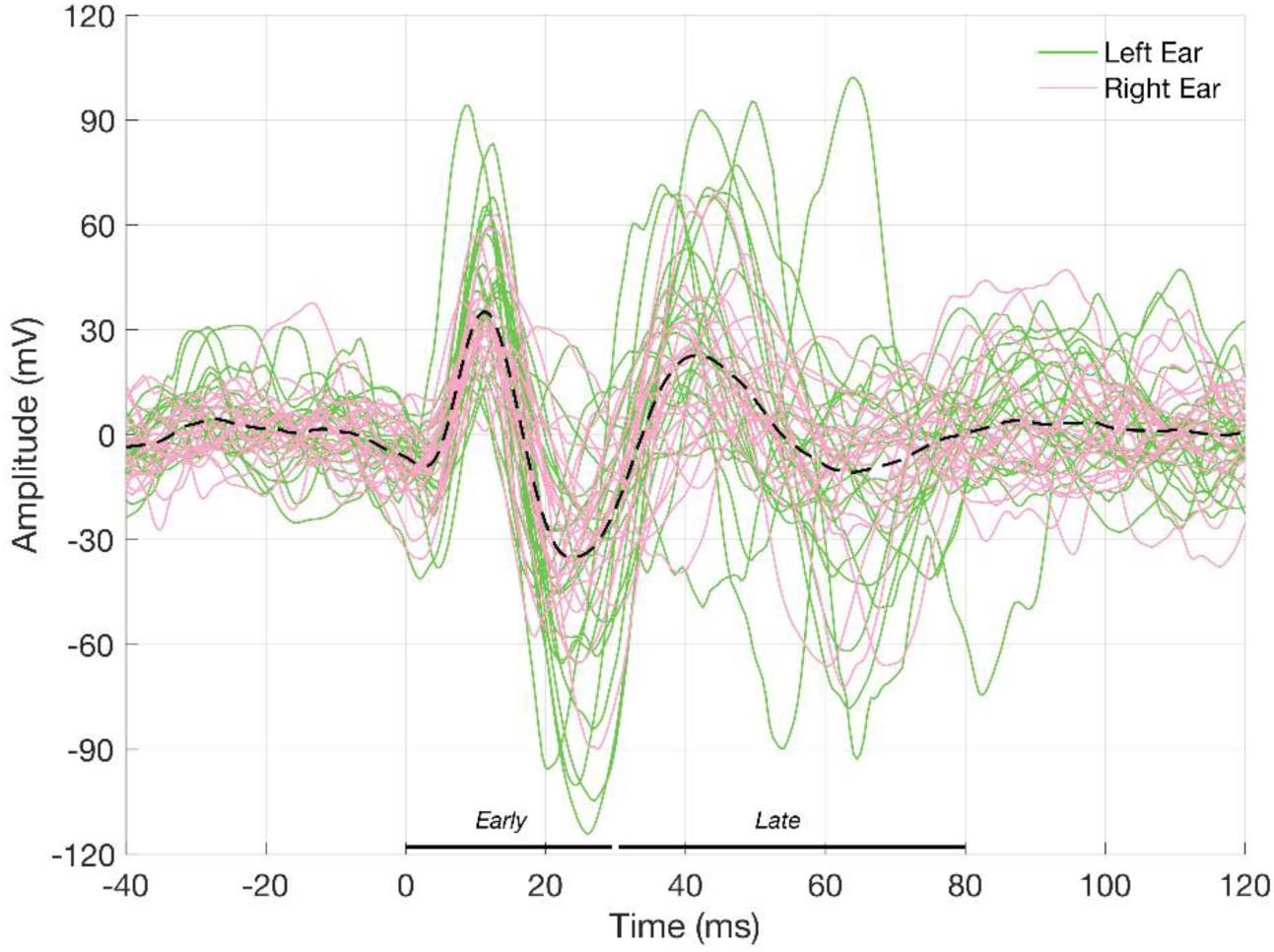
**Individual EMREOs traces**. For each participant the average EMREO was computed separately for the left ear (green) and right ear (pink) and by adjusting the sign to align the phase of EMREOs obtained during ipsi-and contralateral saccades. Each line represents one participant and the dashed line indicates the group mean. The black lines mark the time windows defined as early and late EMREOs.

### 3.3. The EMREO time course does not differ between visually and auditory guided saccades

We then compared EMREOs between paradigms guiding saccades by visual and auditory targets. Because participants found it more difficult to perform auditory guided than visually guided saccades, we obtained considerably fewer valid data epochs for the former paradigm (see Methods and Insets in Figure 5). To boost statistical power, we collapsed EMREOs between ears, adjusting for the phase difference between ipsi-and contralateral EMREOS, which seems valid given the above results. Figure 5 shows the comparison of visual and auditory paradigms separately for each saccade direction and reveals largely similar EMREO time courses. The statistical tests revealed no significant differences between conditions (at p < 0.05; cluster-based permutation statistics).

**Figure 5.**
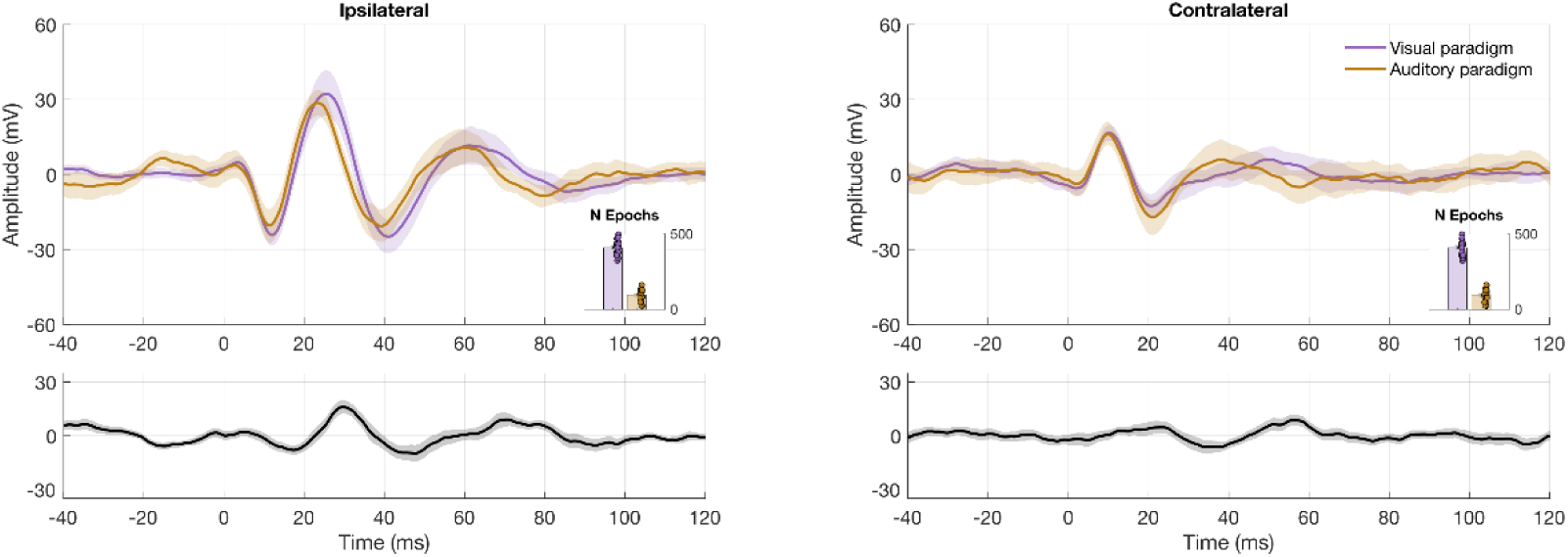
Group-level EMREOs for visually and auditory guided saccades. The panel shows the EMREO time course for saccades towards visual (violet) and auditory (orange) targets. The data points were collapsed across ears after adjusting the sign of ipsi-and contralateral saccades. The lower panels show the difference between paradigms. Lines indicate the group mean (n = 23), shaded areas the 95th percentile bootstrap confidence interval. Insets indicate the number of data epochs per each participant and the group.

### 3.4. Relation between EMREOs, hearing status and middle ear assessments

We tested all participants for normal hearing and middle ear function using pure tone audiometry (PTA) and tympanometric assessment. The audiometric data served as exclusion criteria for the study (see session 2.1), the results from the tympanometric assessment did not. Figure 6 illustrates the pure tone thresholds and thresholds for acoustic reflexes for the included participants. Given that we combined the data across left and right ears to relate the results from ear-assessment in the analysis below, we also tested whether the respective values are comparable between ears. PTA thresholds (averaged across 500, 1000, 2000 and 4000 Hz) for the right (mean ± std = 11.19 ± 4.83 dB SPL) and left ear (9.83 ± 5.76 dB SPL) did not differ significantly (t-test, p = 0.39, t = 0.86). Similarly, the reflexes thresholds for contralateral (right ear: 94.29 ± 6.48 dB SPL; left ear: 98.10 ± 11.01 dB SPL) and ipsilateral stimulation (right ear: 87.84 ± 5.97 dB SPL; left ear: 88.92 ± 4.92 dB SPL) also did not present significant differences (p = 0.17, t = - 1.39 for contralateral reflexes: p = 0.52, t =-0.65 for ipsilateral reflexes).

**Figure 6:**
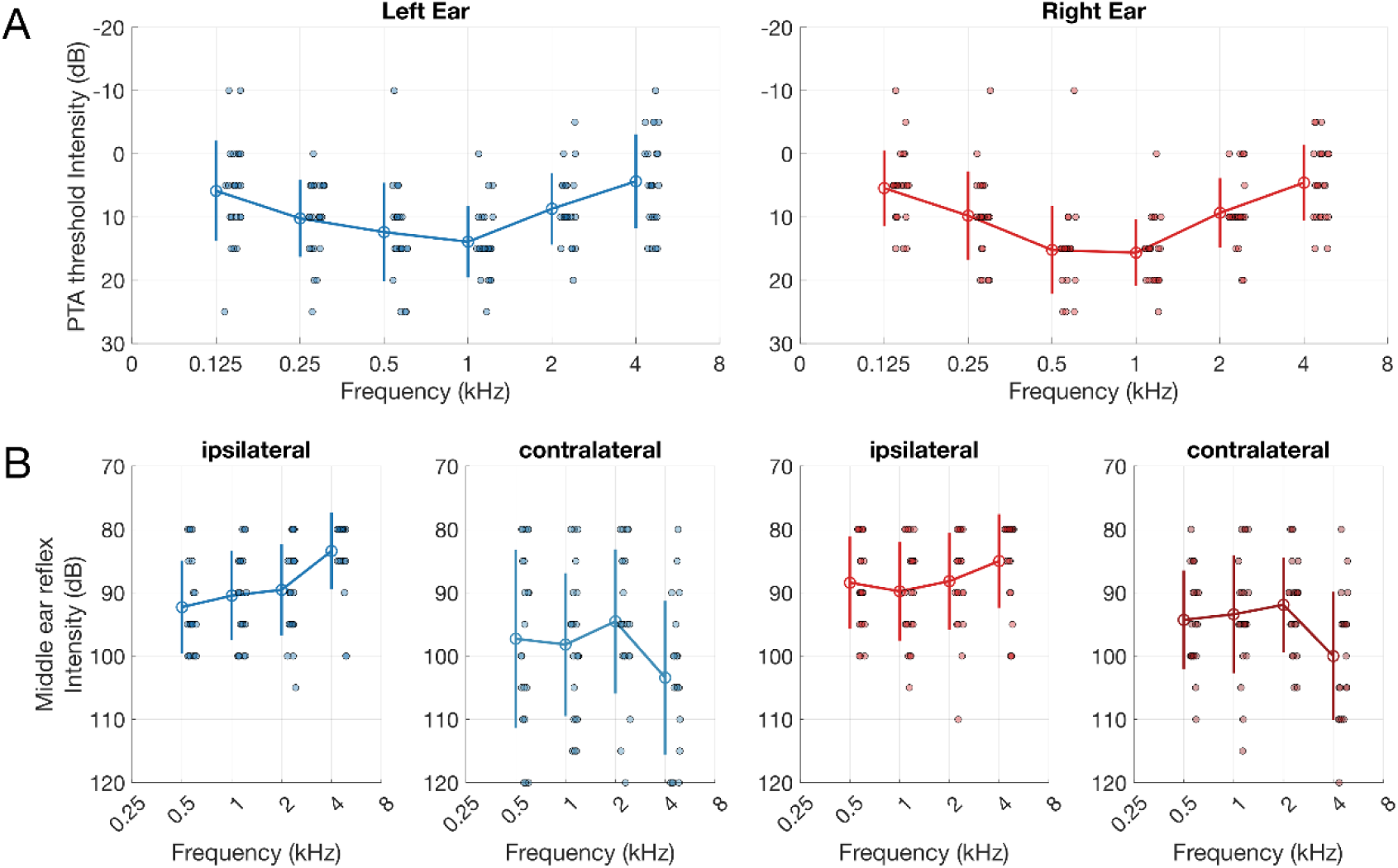
**Pure tone audiometry and middle ear reflexes**. The data shows the individual participant thresholds (dots) and the group averages (lines) for PTA (A) and acoustic reflexes (B). In B, reflexes are divided into ipsi-and contralateral. Blue shades represent left ears and red shades represent right ears. The error bars indicate the group mean and standard deviation (n = 23).

From the tympanometry we also obtained measures of ear canal volume (ECV) and tympanic membrane compliance. For some few individuals (see Fig. 6), these values were slightly outside normal standards (for compliance this would be 0.3 to 1.6 ml according to Jerger, Jerger and Mauldin, 1972): for the present data ECV ranged from 0.36 to 1.88 ml and compliance from 0.25 to 1.28 ml for the right ear, ECV from 0.3 to 1.9 ml and compliance from 0.2 to 1.98 ml for the left ear. However, the group averages of the ECV and compliance were within the expected range for a normal-hearing adult population in both the right (ECV: mean ± std = 0.89 ± 0.49 ml; compliance: 0.62 ± 0.30 ml) and left ear (ECV: 0.80 ± 0.46 ml; compliance: 0.71 ± 0.42 ml). Again we compared the data between ears and found no significant differences (Mann-Whitney test, p = 0.55, U = 568.5 for ECV; p = 0.59, U = 516 for the compliance).

We then asked whether the prominence of an EMREO in an individual was related to these characteristics obtained from the ear assessment. To test this, we computed the overall EMREO amplitude for each ear and participant, defined as the integral of the absolute EMREO deflection (i.e. regardless of sign). The motivation for this definition was that the EMREO time course comprises multiple positive and negative deflections that generally co-occur, and the overall amplitude provides an estimate of the general presence of the EMREO signal. We defined this amplitude both in an early phase of the EMREO (from 0 to 30 ms, c.f. Figure 4) and a later phase (30 ms to 80 ms) and correlated these with the four characteristics. This revealed, as presented in Figure 7, a significant correlation between the early EMREO amplitude and ear canal volume (ECV; rho =-0.45, p = 0.002, n = 46 ears) and eardrum compliance (rho =-0.32, p = 0.03, n = 46 ears), but not with pure tone thresholds (PTA; rho =-0.07, p = 0.61, n = 46 ears) or acoustic reflexes (rho = 0.05, p = 0.76, n = 44 ears). Repeating this analysis for the later EMREO amplitude did not reveal any significant correlations (PTA: rho = - 0.19, p = 0.20; and reflexes: rho =-0.1, p =0.52; ECV: rho =-0.15, p = 0.32; compliance: rho =-0.14, p = 0.35).

**Figure 7.**
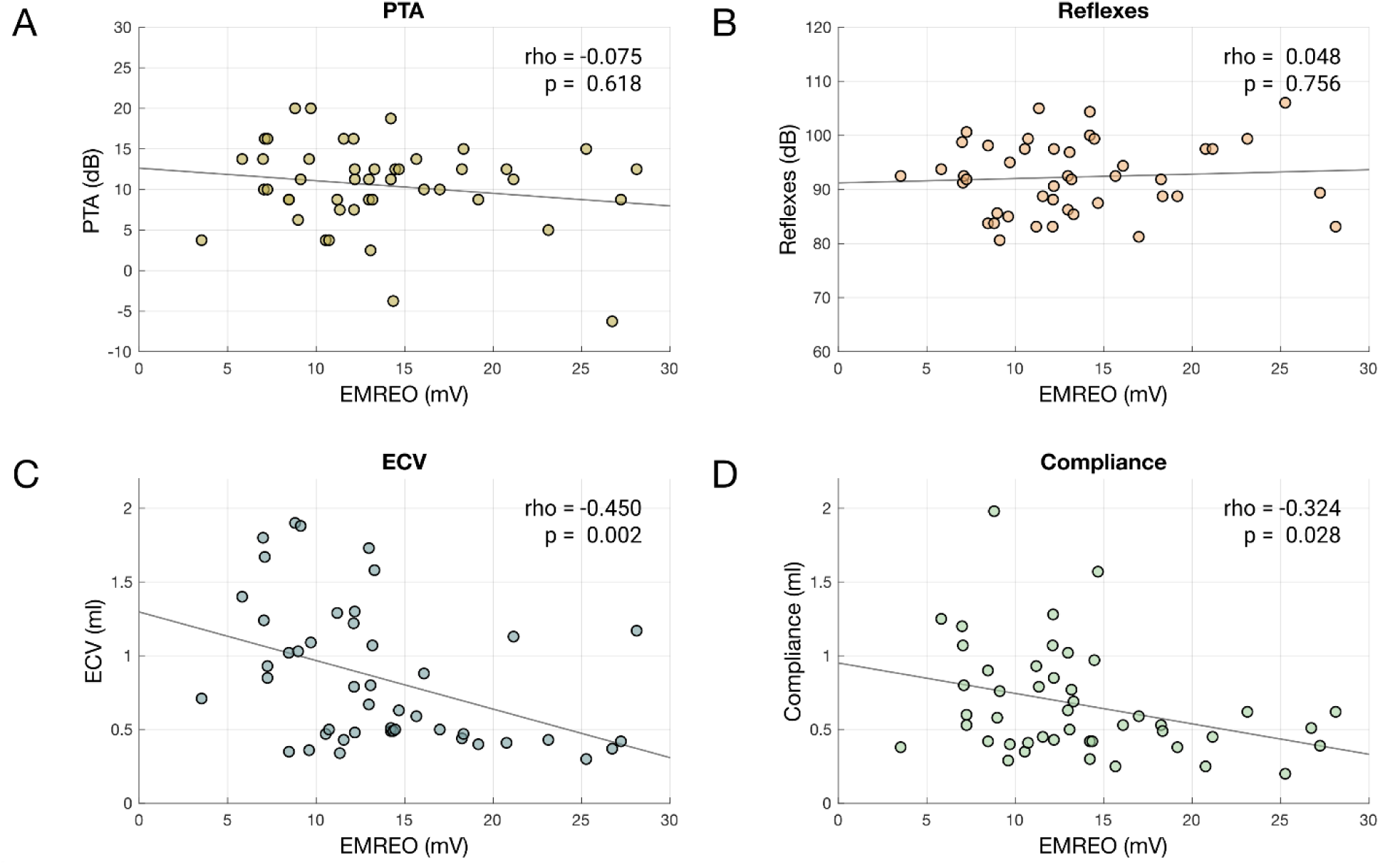
EMREO amplitude in the early phase (0 - 30ms) compared to PTA, tympanometry findings and middle ear reflexes. The scatter plots show the overall EMREO amplitudes (from 0 to 30 ms) against (A) average PTA thresholds, (B) middle ear reflex thresholds across frequencies (mean for ipsi-and contralateral stimulation), (C) equivalent ear canal volume (ECV), and (D) tympanic compliance. Spearman’s correlation rho and p-value are indicated in each panel. Each dot represents one individual ear (n = 46). The grey line shows the linear fit to the data.

## 4. Discussion

Eye movement-related eardrum oscillations have been described in a number of recent studies, though their origin and precise function remain unclear. The overall time course of the EMREOs observed here is qualitatively similar to that reported in previous work and our data confirm the robustness of this phenomenon (Gruters et al., 2018; Lovich et al., 2023a; King et al., 2023; Bröhl and Kayser, 2023). Characteristic features of the EMREO are prominent deflections of latencies around 10 and 20 ms after saccade onset, with an opposing phase in the left and right ears for saccades in the same direction. While previous studies noted that EMREOs look overall similar between left and right ears, those did not provide direct statistical comparison of the moment-by-moment EMREO time course between ears. Based on the comparative analysis of EMREOs bilateral recordings, we here show that indeed the time course and amplitude is comparable between ears. This result is by itself not trivial, given that hemispheric specializations are well known along the auditory pathways (Zatorre, 2022; Jamison et al., 2006; Schönwiesner, Rübsamen and von Cramon, 2005) and studies on otoacoustic emissions have noted potential differences between these between the left and right ears (Keefe et al., 2008; Sininger and Cone-Wesson, 2004).

### 4.1. EMREO amplitude time course differentiates ipsi-and contralateral saccades

The opposing phase of the EMREO between ears for a given saccade, and the opposing phase for ipsi-and contralateral saccades within each ear, have been described in previous studies. Indeed, EMREOs scale both with saccade direction and saccade size, and do so both in horizontal and vertical directions (Murphy et al., 2020; Lovich et al., 2023a). However, previous studies have also noted that the precise EMREO time course may differ between saccades to ipsi-and contralateral targets (Lovich et al., 2023a, King et al., 2023, Bröhl and Kayser, 2023). The present data corroborate this observation and demonstrate statistically significant differences in EMREO amplitude time course for saccades in opposing directions beyond the known opposing phase of these. These differences seem most pronounced starting around 20 ms from saccade onset. Given that the EMREO signal has a characteristic pseudo-rhythmic shape, it is difficult to tell whether this difference reflects a mere scaling of amplitude or a genuine and fundamental difference in the underlying processes.

Noteworthy, this difference in EMREO time course for opposing saccades can provide interesting perspectives on the potential pathways generating the EMREO. Saccades are conjugated eye movements that involve the simultaneous contraction of agonistic and relaxation of antagonistic muscles to facilitate coordinated motion. During ipsilateral saccades, synchronized contraction is mediated by the abducens nucleus, while the contralateral paramedian pontine reticular formation inhibits opposing muscles, demonstrating the distinct neural complexes engaged in relaying differential activation patterns to extraocular muscles (Purves et al., 2001). Similarly, the two middle ear muscles are innervated by different cranial nerves (the fifth for the tensor tympani and the seventh for the stapedius), but there seems to be less knowledge about potential projections from the centers controlling ocular muscles to the pathways controlling the middle ear muscles. Though previous studies have noted that stapedius motor neurons are responsive to non-acoustic sensory signals (Tasko et al., 2022; Deiters et al., 2019; Salomon and Starr, 1963), the pathways and the nature of potential non-acoustic inputs to the middle ear motor neurons remain unclear. Given the opposing activation patterns that control saccadic eye movements, we speculate that the spatial opposition is similarly transferred onto the pathways that innervate middle ear muscles, hence giving rise to distinct EMREO time courses for saccades to ipsi-and contralateral directions.

The observation of different time courses for opposing saccades may also have implications for interpreting the EMREO in the context of aligning spatial stimuli received in eye-and head-centered reference frames (Krüger et al., 2016; Cui et al., 2010; Groh and Sparks, 1992). If the EMREO amplitude would scale in the same manner with saccade amplitude in both saccade directions, reading out such spatial information would be straightforward. However, more fundamental differences in the EMREO time course for opposing saccades would at least require a more sophisticated down-stream readout, calling for further work on how good of a spatial coordinate transformation the EMREO would actually allow.

### 4.2. EMREOs scale with eardrum compliance and ear canal volume

We found that the overall EMREO amplitude in the first 30 ms correlates negatively with the ear canal volume and the compliance of the tympanic membrane. The relation to ear canal volume may simply reflect an acoustic phenomenon whereby a larger canal has a larger acoustic impedance. However, the relation to compliance may be more meaningful. It is possible that some tympanic membranes are under stronger tension from the middle ear muscles and thereby show a lower compliance. If the EMREO is indeed generated by the middle ear muscles, stronger tension could indicate greater control of the membrane by those pathways. Hence, while some ears may exhibit a stronger influence from the pathways driving EMREOs and thus reduced flexibility, others may show the opposite. Alternatively, the differences in compliance might be due to a more rigid ossicular chain. This could passively lead to larger amplitudes if the tympanic membrane is more sensitive to ringing at eigenfrequencies closely related to that of EMREOs, or in larger amplitudes if the underlying process includes an active overcompensation to counteract an otherwise less variable sound transduction. Still, it is not immediately clear why in such a case one would observe a significant correlation of compliance only with the early but not the late EMREO phase.

One possibility is that the earlier phase of the EMREO is more directly related to the actual phenomenon generating the EMREO while the later phase reflects simply the ringing of the membrane in response to an abrupt stiffening or relaxation. It could be that the damped oscillatory signal that one obtains with in-ear microphones directly corresponds to an oscillatory drive exerted on the tympanic membrane. Alternatively, it could be that the tympanic membrane simply rings in response to an abrupt stiffening or relaxation induced by changes in the tension of a particular muscle, as theorized by Cho and colleagues (2023). More work is required to investigate these possibilities.

It has been speculated that EMREOs result from motor element activity of the auditory system, namely middle ear muscles and outer hair cells. Both can be modulated by signals descending from the cortex and higher structures of the auditory pathway (Gruters et al., 2018; Gallagher, Diop and Olson, 2021; Lovich et al., 2023a). Here we tested variables related to the functioning of the middle ear muscles and properties of the ear canal and eardrum, but did not specifically address the role of outer hair cells in the generation of EMREOs, leaving the precise contribution of the latter unclear.

### 4.3. EMREOs relate to motor but not sensory signals

Our data also show that the EMREO is comparable when saccades are made towards a visual target to when saccades are guided by a sound. This fits with previous work reporting EMREOs obtained during spontaneous viewing of images or in darkness (Lovich et al., 2023b). Overall, this suggests that the EMREO is mostly related to the motor act of a saccade and not exogenous sensory information. We note that participants had considerable difficulties performing purely acoustically guided saccades towards targets, which resulted in very different numbers of data epochs in both conditions. While this leaves the comparison of auditory and visually-guided EMREOs dependent on unequal numbers of data epochs, we do not believe that this critically influences the statistical result obtained here. The conclusion that the EMREO is mostly related to the motor action of the saccade rather than a sensory cue is commensurate with previous results, and is also supported by our previous study where we showed that the presence of an attentional cue did not systematically alter the EMREO (Bröhl and Kayser, 2023).

The motor link between eye movements and the EMREOs relates to a larger topic of extrinsic auricular muscle synkinesis in relation to gaze movements (Wilson, 1908; Carmichael and Critchley, 1924, 1925; Schmidt and Thoden, 1978). Developmentally, the cranial nerves controlling eye movements, the middle ear muscles and those controlling the auricular muscles all share the same mesoblastic origin (Carmichael and Critchley, 1925). The circuits activating the auricular muscles overlap with those most likely giving rise to the EMREO. The postauricular reflex, wherein the posterior auricular muscle activates in response to loud sounds, is supposedly guided by acoustic signals related to the facial motor nucleus from the ascending auditory pathways via the lateral lemniscus, but also indirectly via the reticular nucleus from the superior colliculus (Liugan, Zhang and Cakmak, 2018; O’Beirne and Patuzzi., 1999; Yoshie and Okudaira, 1969). The latter opens the pathway for signals relating to eye movements but also visual stimuli to activate auricular and possibly also middle ear muscles. While future work needs to define the precise contributions of the two middle ear muscles to the EMREO and the pathways controlling the EMREO, the broader topic of how non-acoustic information influences the functioning of the muscles in and around the ear remains exciting.

## REFERENCES

1. Bon, L., & Lucchetti, C. (1988). The motor programs of monkey’s saccades: an attentional hypothesis. Experimental brain research, 71, 199–207. 10.1007/BF00247535

2. Borg, E. R. I. K., Counter, S. A., & Rösler, G. (1984). Theories of middle-ear muscle function. The acoustic reflex: Basic principles and clinical applications, 63-99.

3. Brainard, D. H. (1997). The Psychophysics Toolbox. Spatial Vision, 10(4), 433–436. 10.1163/156856897X00357

4. Bröhl, F., & Kayser, C. (2023) Detection of Spatially Localized Sounds Is Robust to Saccades and Concurrent Eye Movement-Related Eardrum Oscillations (EMREOs). Journal of Neuroscience, 43 (45) 7668–7677; 10.1523/JNEUROSCI.0818-23.2023

5. Carmichael, E. A., & Critchley, M. (1924). Facial associated movements. Journal of Neurology and Psychopathology, 5(18), 124. 10.1136/jnnp.s1-5.18.124

6. Carmichael, E. A., & Critchley, M. (1925). The relations between eye movements and other cranial muscles. The British Journal of Ophthalmology, 9(2), 49. 10.1136/bjo.9.2.49

7. Caruso, V. C., Pages, D. S., Sommer, M. A., & Groh, J. M. (2021). Compensating for a shifting world: evolving reference frames of visual and auditory signals across three multimodal brain areas. Journal of neurophysiology, 126(1), 82–94. https://journals.physiology.org/doi/full/10.1152/jn.00385.2020

8. Cho, N. H., Ravicz, M. E., & Puria, S. (2023). Human middle-ear muscle pulls change tympanic-membrane shape and low-frequency middle-ear transmission magnitudes and delays. Hearing Research, 430, 108721.. 10.1016/j.heares.2023.108721

9. Cui, Q. N., Razavi, B., O’Neill, W. E., & Paige, G. D. (2010). Perception of auditory, visual, and egocentric spatial alignment adapts differently to changes in eye position. Journal of neurophysiology, 103(2), 1020–1035. 10.1152/jn.00500.2009

10. Deiters, K. K., Flamme, G. A., Tasko, S. M., Murphy, W. J., Greene, N. T., Jones, H. G., & Ahroon, W. A. (2019). Generalizability of clinically measured acoustic reflexes to brief sounds. The Journal of the Acoustical Society of America, 146(5), 3993. 10.1121/1.5132705

11. Edmonson, A., Iwanaga, J., Olewnik, Ł., Dumont, A. S., & Tubbs, R. S. (2022). The function of the tensor tympani muscle: a comprehensive review of the literature. Anatomy & Cell Biology, 55(2), 113–117. 10.5115/acb.21.032

12. Gallagher, L., Diop, M., & Olson, E. S. (2021). Time-domain and frequency-domain effects of tensor tympani contraction on middle ear sound transmission in gerbil. Hearing research, 405, 108231. 10.1016/j.heares.2021.108231

13. Groh, J. M., & Sparks, D. L. (1992). Two models for transforming auditory signals from head-centered to eye-centered coordinates. Biological cybernetics, 67(4), 291–302. 10.1007/BF02414885

14. Gruters, K. G., Murphy, D. L., Jenson, C. D., Smith, D. W., Shera, C. A., & Groh, J. M. (2018). The eardrums move when the eyes move: A multisensory effect on the mechanics of hearing. Proceedings of the National Academy of Sciences, 115(6), E1309–E1318. 10.1073/pnas.1717948115

15. Jamison, H. L., Watkins, K. E., Bishop, D. V., & Matthews, P. M. (2006). Hemispheric specialization for processing auditory nonspeech stimuli. Cerebral cortex, 16(9), 1266–1275. 10.1093/cercor/bhj068

16. Jerger, J., Jerger, S., & Mauldin, L. (1972). Studies in impedance audiometry: I. Normal and sensorineural ears. Archives of Otolaryngology, 96(6), 513–523. 10.1001/archotol.1972.00770090791004

17. Keefe, D. H., Gorga, M. P., Jesteadt, W., & Smith, L. M. (2008). Ear asymmetries in middle-ear, cochlear, and brainstem responses in human infants. The Journal of the Acoustical Society of America, 123(3), 1504–1512. 10.1121/1.2832615

18. King, C. D., Lovich, S. N., Murphy, D. L. K., Landrum, R., Kaylie, D., Shera, C. A., & Groh, J.M. (2023). Individual similarities and differences in eye-movement-related eardrum oscillations (EMREOs). Hearing Research, 440, 108899. 10.1016/j.heares.2023.108899

19. Krüger, H. M., Collins, T., Englitz, B., & Cavanagh, P. (2016). Saccades create similar mislocalizations in visual and auditory space. Journal of neurophysiology, 115(4), 2237–2245. 10.1152/jn.00853.2014

20. Leigh, R. J., & Kennard, C. (2004). Using saccades as a research tool in the clinical neurosciences. Brain, 127(3), 460–477. 10.1093/brain/awh035

21. Liugan, M., Zhang, M., & Cakmak, Y. O. (2018). Neuroprosthetics for auricular muscles: neural networks and clinical aspects. Frontiers in neurology, 8, 752. 10.3389/fneur.2017.00752

22. Lovich, S. N., King, C. D., Murphy, D. L., Landrum, R. E., Shera, C. A., & Groh, J. M. (2023a). Parametric information about eye movements is sent to the ears. Proceedings of the National Academy of Sciences, 120(48), e2303562120. 10.1073/pnas.2303562120

23. Lovich, S. N., King, C. D., Murphy, D. L., Abbasi, H., Bruns, P., Shera, C. A., & Groh, J. M. (2023b). Conserved features of eye movement related eardrum oscillations (EMREOs) across humans and monkeys. Philosophical Transactions of the Royal Society B, 378(1886), 20220340. 10.1098/rstb.2022.0340

24. Lucchetti, C., & Bon, L. (1997). Motor programs of spontaneous and visually guided saccades in macaque monkey: An electrophysiological approach. International journal of neuroscience, 90(1-2), 37–43.. 10.3109/00207459709000624

25. Machado, A.B.M. (2004). *Functional Neuroanatomy*. 2nd edition, Atheneu.

26. Maris, E., & Oostenveld, R. (2007). Nonparametric statistical testing of EEG-and MEG-data. Journal of neuroscience methods, 164(1), 177–190. 10.1016/j.jneumeth.2007.03.024

27. Møller, A. R. (1984). Neurophysiological basis of the acoustic middle-ear reflex. In The acoustic reflex (pp. 1-34). Academic Press.

28. Mukerji, S., Windsor, A. M., & Lee, D. J. (2010). Auditory brainstem circuits that mediate the middle ear muscle reflex. Trends in amplification, 14(3), 170–191. 10.1177/1084713810381771

29. Murphy, D. L., King, C. D., Lovich, S. N., Landrum, R. E., Shera, C. A., & Groh, J. M. (2020). Evidence for a system in the auditory periphery that may contribute to linking sounds and images in space. bioRxiv, 2020-07. 10.1101/2020.07.19.210864

30. Nichols, T. E., & Holmes, A. P. (2002). Nonparametric permutation tests for functional neuroimaging: a primer with examples. Human brain mapping, 15(1), 1–25. 10.1002/hbm.1058

31. O’Beirne, G. A., & Patuzzi, R. B. (1999). Basic properties of the sound-evoked post-auricular muscle response (PAMR). Hearing research, 138(1-2), 115–132. 10.1016/S0378-5955(99)00159-8

32. Pierrot-Deseilligny, C. (2011). Nuclear, internuclear, and supranuclear ocular motor disorders. Handbook of clinical neurology, 102, 319–331. 10.1016/B978-0-444-52903-9.00018-2

33. Purves, D., Augustine, G., Fitzpatrick, D., Katz, L., LaMantia, A., McNamara, J., & Williams, S. (2001). Neuroscience 2nd edition. sunderland (ma) sinauer associates. Types of Eye Movements and Their Functions.

34. Razavi, B., O’Neill, W. E., & Paige, G. D. (2007). Auditory spatial perception dynamically realigns with changing eye position. Journal of Neuroscience, 27(38), 10249–10258. 10.1523/JNEUROSCI.0938-07.2007

35. Salomon, G., & Starr, A. (1963). Electromyography of middle ear muscles in man during motor activities. Acta Neurologica Scandinavica, 39(2), 161–168. 10.1111/j.1600-0404.1963.tb05317.x

36. Schmidt, D., & Thoden, U. (1978). Co-activation of the M. transversus auris with eye movements (Wilson’s oculo-auricular phenomenon) and with activity in other cranial nerves. Albrecht von Graefes Archiv für klinische und experimentelle Ophthalmologie, 206, 227–236. 10.1007/BF02387334

37. Schönwiesner, M., Rübsamen, R. & Von Cramon, D.Y. (2005), Hemispheric asymmetry for spectral and temporal processing in the human antero-lateral auditory belt cortex. European Journal of Neuroscience, 22: 1521–1528. 10.1111/j.1460-9568.2005.04315.x

38. Sininger, Y. S., & Cone-Wesson, B. (2004). Asymmetric cochlear processing mimics hemispheric specialization. Science, 305(5690), 1581–1581. 10.1126/science.1100646

39. Tasko, S. M., Deiters, K. K., Flamme, G. A., Smith, M. V., Murphy, W. J., Jones, H. G.,… & Ahroon, W. A. (2022). Effects of unilateral eye closure on middle ear muscle contractions. Hearing research, 424, 108594. 10.1016/j.heares.2022.108594

40. The MathWorks Inc. (2022). MATLAB version: 9.13.0 (R2022b), Natick, Massachusetts: The MathWorks Inc. https://www.mathworks.com

41. Walker, H. K., Hall, W. D., & Hurst, J. W. (1990). Cranial nerves III, IV, and VI: the oculomotor, trochlear, and abducens nerves. In Clinical Methods: The History, Physical, and Laboratory Examinations. 3rd edition. Butterworths.

42. Wilson, S.A.K. (1908). A note on an associated movement of the eyes and ears in man. Review of Neurology and Psychiatry, 6, 331.

43. World Health Organization. (2021). World report on hearing. https://www.who.int/publications/i/item/9789240020481

44. Yoshie, N., & Okudaira, T. (1969). Myogenic evoked potential responses to clicks in man. Acta oto-laryngologica, 67(sup252), 89–103. 10.3109/00016486909120515

45. Zatorre, R. J. (2022). Hemispheric asymmetries for music and speech: Spectrotemporal modulations and top-down influences. Frontiers in neuroscience, 16, 1075511.

